# Metagenomic Insights into Microbial Metabolisms of a Sulfur-Influenced Glacial Ecosystem

**DOI:** 10.1101/2020.01.31.929786

**Authors:** Christopher B. Trivedi, Blake W. Stamps, Graham E. Lau, Stephen E. Grasby, Alexis S. Templeton, John R. Spear

**Affiliations:** Department of Civil and Environmental Engineering, Colorado School of Mines, Golden, CO, 80401 USA; Department of Geological Sciences, University of Colorado Boulder, Boulder, CO, 80309 USA; Geological Survey of Canada-Calgary, Calgary, AB, T2L2A7 Canada

## Abstract

Biological sulfur cycling in polar, low-temperature ecosystems is an understudied phenomenon in part due to difficulty of access and the ephemeral nature of such environments. One such environment where sulfur cycling plays an important role in microbial metabolisms is located at Borup Fiord Pass (BFP) in the Canadian High Arctic. Here, transient springs emerge from the toe of a glacier creating a large proglacial aufeis (spring-derived ices) that are often covered in bright yellow/white sulfur, sulfate, and carbonate mineral precipitates that are accompanied by a strong odor of hydrogen sulfide. Metagenomic sequencing from multiple sample types at sites across the BFP glacial system produced 31 highly complete metagenome assembled genomes (MAGs) that were queried for sulfur-, nitrogen- and carbon-cycling/metabolism genes. Sulfur cycling, especially within the Sox complex of enzymes, was widespread across the isolated MAGs and taxonomically associated with the bacterial classes *Alpha-, Beta-, Gamma-*, and *Epsilon- Proteobacteria*. While this does agree with previous research from BFP implicating organisms within the *Gamma-* and *Epsilon- Proteobacteria* as the primary classes responsible for sulfur oxidation, our new data suggests putative sulfur oxidation by organisms within *Alpha-* and *Beta- Proteobacterial* classes which was not predicted. These findings indicate that in a low-temperature, ephemeral sulfur-based environment such as this, functional redundancy may be a key mechanism that microorganisms use to co-exist whenever energy is limited and/or focused by redox chemistry.

**Importance:** Borup Fiord Pass is a unique environment characterized by a sulfur-enriched glacial ecosystem, in the low-temperature environment of the Canadian High Arctic. This unique combination makes BFP one of the best analog sites for studying icy, sulfur-rich worlds outside of our own, such as Europa and Mars. The site also allows investigation of sulfur-based microbial metabolisms in cold environments here on Earth. Herein, we report whole genome sequencing data that suggests sulfur cycling metabolisms at BFP are more widely used across bacterial taxa than predicted. From our data, the metabolic capability of sulfur oxidation among multiple community members appears likely due to functional redundancy within their genomes. Functional redundancy, with respect to sulfur-oxidation at BFP, may indicate that this dynamic ecosystem hosts microorganisms that are able to use multiple sulfur electron donors alongside other important metabolic pathways, including those for carbon and nitrogen.

## 1 Introduction

Arctic and Antarctic ecosystems such as lakes (Jungblut et al., 2016), streams (Niederberger et al., 2009; Andriuzzi et al., 2018), groundwater-derived ice (aufeis; Grasby et al., 2014; Hodgkins et al., 2004), and saturated sediments (Lamarche-Gagnon et al., 2015) provide insight into geochemical cycling and microbial community dynamics in low temperature environments. One such ecosystem in the Canadian High Arctic, Borup Fiord Pass (BFP), is located on northern Ellesmere Island, Nunavut, Canada and contains a transient sulfidic spring system discharging through glacial ice and forming supra and proglacial ice deposits (aufeis) as well as cryogenic mineral precipitates (Figure 1; Grasby et al., 2003).

**Figure 1.**
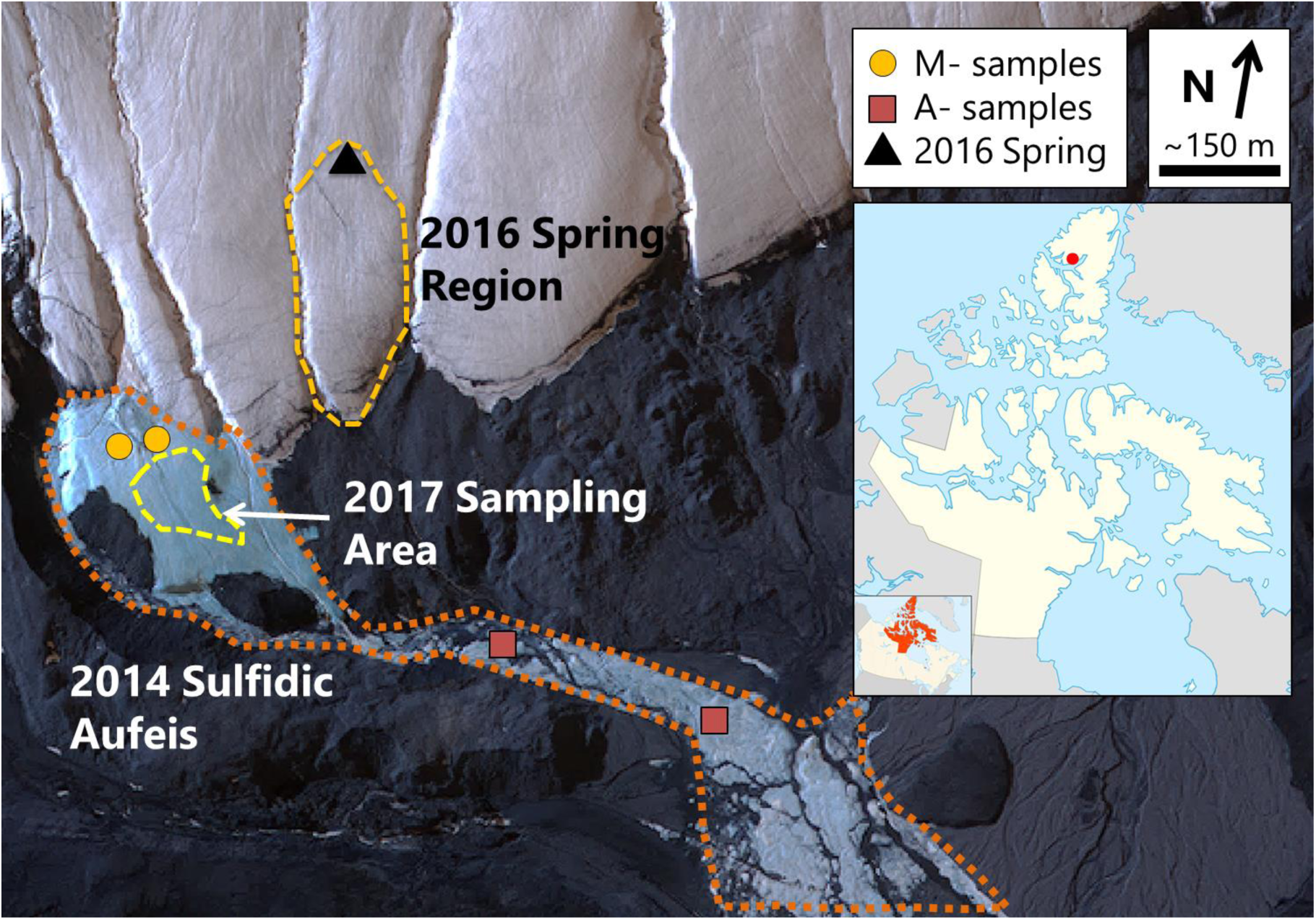
Borup Fiord Pass satellite view. Image was collected on 2 July 2014. M- and A-samples from 2014 are marked with gold circles and red squares, respectively. The 2016 Spring region is highlighted with a dashed gold line, and the location of the spring is marked with a black triangle. Samples from 2017 were collected from within yellow dashed area. The inset line map shows the approximate location of Borup Fiord Pass (red dot) on Ellesmere Island, Nunavut, Canada. *Satellite image courtesy of the DigitalGlobe Foundation, and inset line map courtesy of Wikimedia Commons and the CC 3*.*0 license*.

Borup Fiord Pass is a north-south trending valley through the Krieger mountains at 81°01’ N, 81°38’ W (Figure 1, inset). The spring system of interest is approximately 210-240 m above sea level and occurs at the toe of two coalesced glaciers (Grasby et al., 2003). The spring system is thought to be perennial in nature but discharges from different locations along the glacier from year to year (Grasby et al., 2003; Gleeson et al., 2010; Wright et al., 2013; Lau et al., 2017). However, given lack of winter observations it is not confirmed that spring flow is year-round or limited to the warmer spring and summer months. However, growth of large aufeis formations through the dark season support continued discharge through the winter. A lateral fault approximately 100 m south of the toe of the glacier has been implicated in playing a role in the subsurface hydrology, in that this may be one of the areas where subsurface fluid flow is focused to the surface (Grasby et al., 2003; Scheidegger, et al. 2012). Furthermore, organic matter in shales along this fault may be a source of carbon for subsurface microbial processes such as biological sulfate reduction (BSR; Lau et al., 2017).

Previous research at BFP has surveyed microbial activity, redox bioenergetics (Wright et al., 2013), the biomineralization of elemental sulfur (S^0^; (Grasby et al., 2003; Gleeson et al., 2011; Lau et al., 2017), and the cryogenic carbonate vaterite (Grasby, 2003). Wright et al. (2013) produced a metagenome in the context of geochemical data to constrain the bioenergetics of microbial metabolism from a mound of elemental sulfur sampled at the site in 2012, and found that at least in surface mineral deposits, sulfur oxidation was likely the dominant metabolism present among *Epsilonproteobacteria*. An exhaustive 16S rRNA gene sequencing study conducted on samples collected in 2014-2017 revealed an active and diverse assemblage of autotrophic and heterotrophic microorganisms in melt pools, aufeis, spring fluid, and surface mineral deposits that persist over multiple years, contributing to a basal community present in the system regardless of site or material type (Trivedi et al., 2018). However, to date no work has been attempted to fully understand and characterize the metabolic capability of the microbial communities adapted to this glacial ecosystem, including sulfur metabolism beyond the surface precipitates at the site.

Microbial sulfur oxidation and reduction, which spans an eight electron transfer between the most oxidized and reduced states, is important in a multitude of diverse extreme environments, such as hydrothermal vents & deep subsurface sediments (Dahl and Friedrich, 2008), microbial mats (Ley et al., 2006; Feazel et al., 2008; Robertson et al., 2009), sea ice (Boetius et al., 2015), glacial environments (Mikucki and Priscu, 2007; Purcell et al., 2014), and Arctic hypersaline springs (Perreault et al., 2008; Niederberger et al., 2009). Two predominant forms of sulfur-utilizing microorganisms exist: sulfur-oxidizing microorganisms (SOMs), which use reduced compounds (e.g. H_2_S and S^0^) as electron donors, and sulfate-reducing microorganisms (SRMs), which use oxidized forms of sulfur (e.g. SO_4_^2-^ and SO_3_^2-^) as electron acceptors. SOMs are metabolically and phylogenetically diverse (Friedrich et al., 2001, 2005; Dahl and Friedrich, 2008), and can often autotrophically fix carbon dioxide (CO_2_) while using a variety of electron acceptors (Mattes et al., 2013). Likewise, SRMs can utilize a range of electron donors, including organic carbon, inorganic carbon, methane, hydrogen, metallic iron (Enning et al., 2012), and even heavier metals such as uranium, where the soluble U(VI) is converted to the insoluble U(IV) under anoxic conditions (Spear et al., 1999, 2000). SRMs are particularly important in sulfate-rich marine environments where they contribute as much as 50% of total global organic carbon oxidation (Thullner et al., 2009; Bowles et al., 2014).

Some research has been conducted on microbial sulfur-cycling in low-temperature polar environments in both Antarctica (Mikucki et al., 2009, 2016) and the Arctic (Niederberger et al., 2009; Purcell et al., 2014), including BFP (Gleeson et al., 2011; Grasby et al., 2012; Wright et al., 2013). However, the organisms that participate in sulfur cycling and which metabolic pathways they utilize in these low-temperature environments remain understudied. It is important that we better classify these systems as they are key to our understanding of how life can adapt to potentially adverse conditions (e.g., low-temperature and sulfidic conditions). Furthermore, these adaptations may also be able to inform us about what to search for on worlds outside of Earth, for instance Europa, where we know that low-temperature, sulfur-rich conditions exist (Carlson et al., 1999), or Mars, where gypsum was discovered in the North Polar Cap (Massé et al., 2012).

We collected samples over multiple years (2014, 2016, and 2017; Figure 2) and from varied sample types (e.g., aufeis, melt pools, spring fluids, and surface mineral precipitates; Figure S2) for metagenomic whole genome sequencing (WGS). These data were assembled and binned into Metagenome Assembled Genomes (MAGs), which were queried to identify which metabolic processes are present and dominant across multiple sample types at BFP. Analysis of these MAGs revealed the presence of sulfur oxidation genes (namely those part of the Sox operon) across multiple taxonomic genomes, including in genomes related to organisms where it is not predicted to be metabolically viable. This may indicate a form of functional redundancy present in the BFP system where organisms from other Phyla are able to take advantage of the abundance of reduced sulfur for metabolic processes. While sulfur metabolism was the focus of this study, genes for carbon, nitrogen, oxygen, hydrogen, and methane metabolisms were queried as this can provide information about energy utilization and potential electron acceptors.

**Figure 2.**
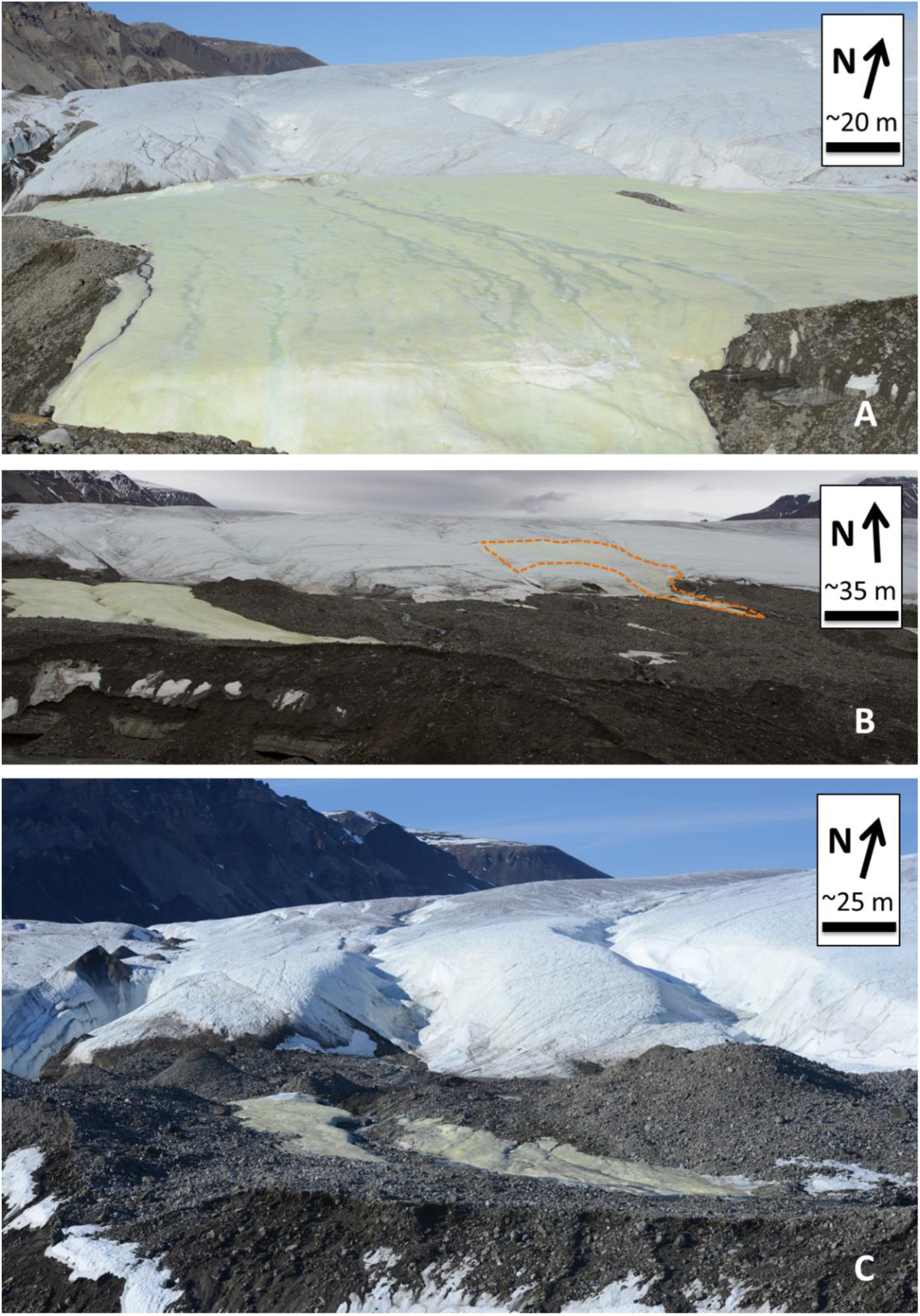
BFP sampling locations. **A)** The “Sulfidic Aufeis” from Trivedi, et al. (2018) during 2014. Note: Two samples locations are to the southwest of this picture and their approximate locations can be seen in Figure 1. **B)** The sulfidic aufeis on the left of the picture with the 2016 Spring a few hundred meters to the east (indicated by the orange dashed line; also see Figure S1). **C)** The same location and focus of the sampling in 2017.

## 2 Results

### Assembly and Identification of Metagenome Assembled Genomes

High throughput sequencing of total environmental genomic DNA (gDNA) extracted from nine BFP biomass samples produced paired-end reads that were trimmed and co-assembled *de novo*. The co-assembly produced 1.16 Gbp in 477,394 contigs, with the largest being 406,785 bp long and the average being 2,440 bp long. A summary of all assembly statistics is available within supplementary Table S1. After assembly, a total of 166 bins were identified after CONCOCT binning (Alneberg et al., 2014) and further refinement within Anvi’o (Eren et al., 2015). Of these, 31 were classified as MAGs (defined as bins with greater than 50% estimated genome completeness and less than 10% redundancy) after quality control within CheckM (Parks et al., 2015). Of the 31 MAGs, nine were considered highly complete with low redundancy (90% and 10%, respectively). Further summary statistics of sequencing data are available in Table 1. MAG distribution was not even between samples. One broadly distributed MAG, MAG 31 (most closely related to the genus *Flavobacterium*) had abundant contribution from samples A6 (aufeis), 3B, 4E, and 6B (all mineral precipitate samples), and moderate contributions from A4b and M4b (aufeis and melt pool samples, respectively). However, MAG 31 had the lowest completion of reported MAGs.

**Table 1.**
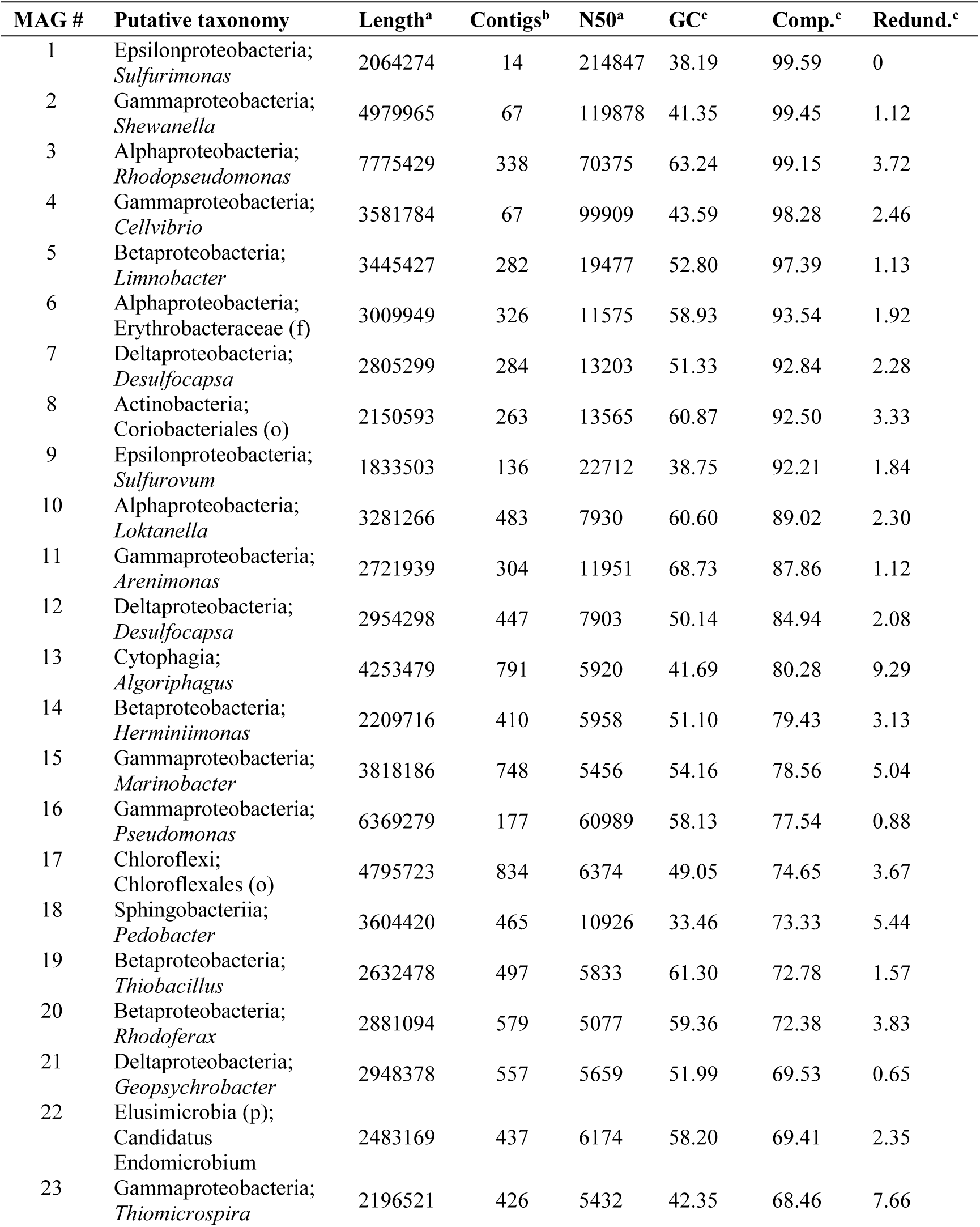
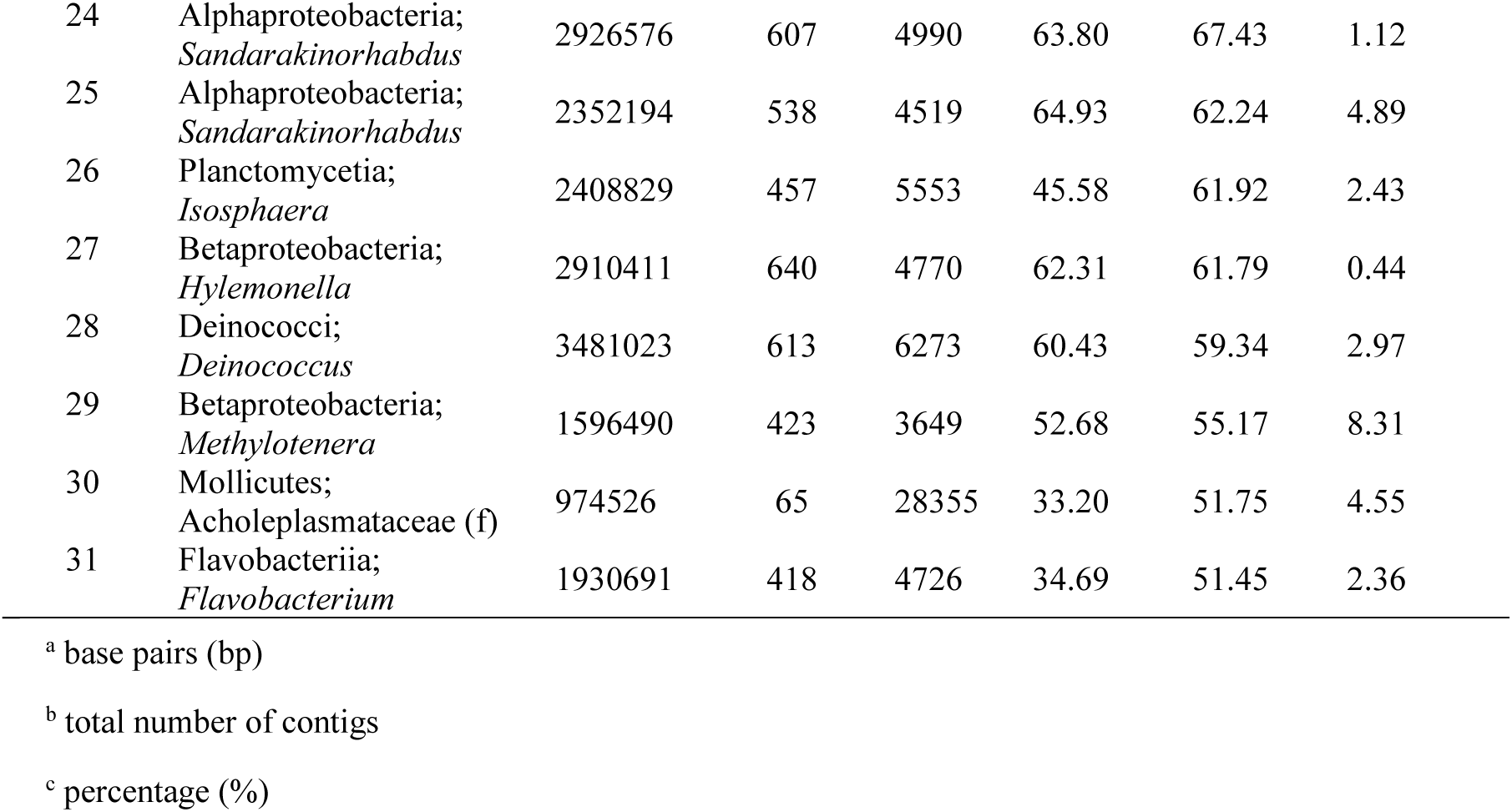
Metadata for all reported MAGs. All MAGs shown are >50% completion with <10% redundancy via CheckM. Nearest neighbors were placed in a cladogram via CheckM/pplacer and confirmed via RAST (Figure 5).

All MAGs were identified within the domain Bacteria. The majority of MAGs (22 of 31) classified within the phylum *Proteobacteria* (Table 1). Of the *Proteobacteria*, the *Betaproteobacteria* and *Gammaproteobacteria* were the most represented, with six (MAG 29, 19, 14, 5, 20, & 27) and six (MAG 11, 23, 2, 4, 15, & 16) MAGs, respectively. The class *Alphaproteobacteria* contained five MAGs (MAG 10, 3, 6, 25, & 25), with the *Deltaproteobacteria* (three; MAG 7, 12, & 21) and *Epsilonproteobacteria* (two; MAG 1 and 9) having the fewest. Two MAGs (7 and 12) were putatively identified by average nucleotide identity (ANI) within the *Deltaproteobacterial* genera *Desulfocapsa* (Table S2). Each class contained at least one highly complete, low redundancy MAG (Table 1).

### Non-Sulfur Associated Metabolic Potential Identified Across BFP

Carbon fixation potential was inferred by looking at three pathways: Wood-Ljundahl (reductive acetyl-coenzyme A); Reverse tricarboxylic acid cycle (rTCA); and ribulose-1,5-bisphosphate carboxylase pathway (RuBisCO). ATP citrate lyase (*acl*), a key enzyme in the rTCA cycle (Perner et al., 2007), was identified in two MAGs (1 and 9 respectively), while genes for other rTCA enzymes were found within 17 of 31 of MAGs (Figure 3b). ATP citrate lyase was found in both *Epsilonproteobacterial* MAGs (*Sulfurimonas* (MAG 1) and *Sulfurovum* (MAG 9)). The most commonly identified carbon fixation pathway within the queried genomes from BFP was the Wood–Ljungdahl pathway found in all but three MAGs (14, 22, and 31). RuBisCo was also identified in four MAGs with both *cbbL* or *cbbM* genes present (10, 19, 23, and 27).

**Figure 3.**
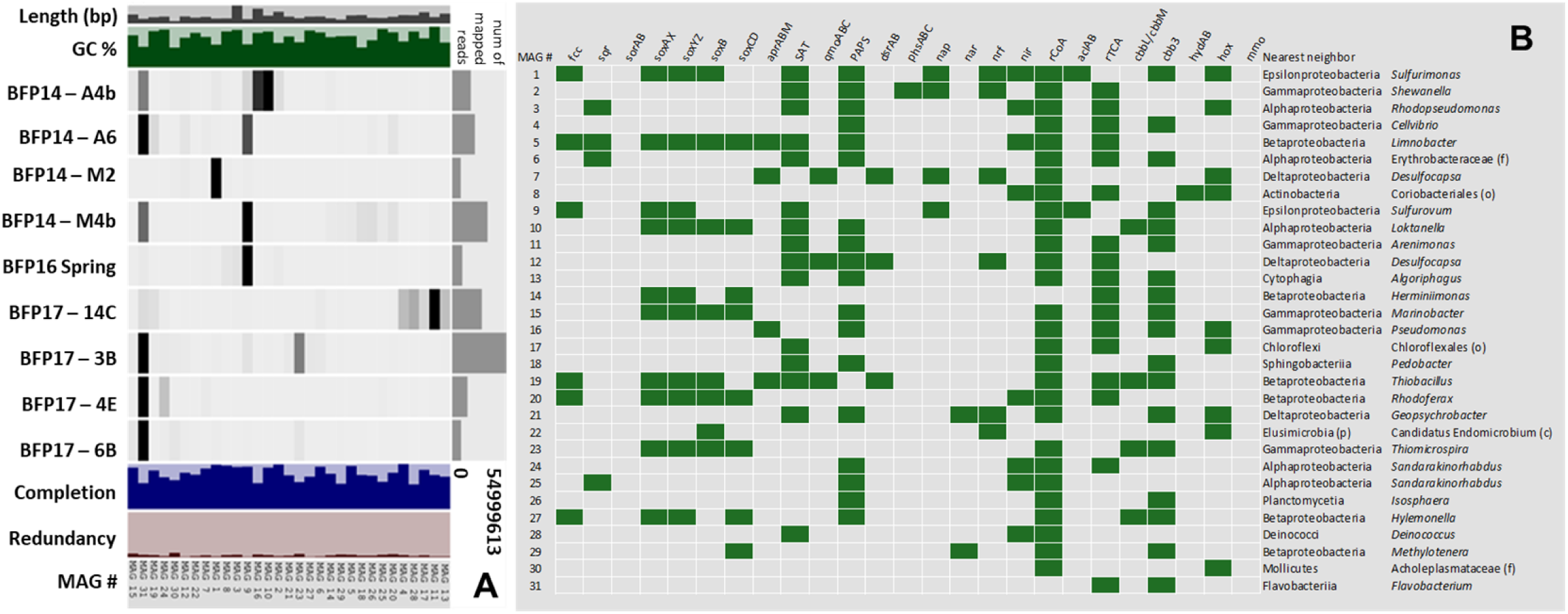
Anvi’o diagram and gene presence/absence table. **A)** This diagram shows all MAGs that are >50% completion and <10% redundancy. Higher contribution to a given MAG by a sample is indicated by darker bars, while lower contribution is indicated by lighter/no bars. **B)** This table shows the presence/absence of the specific genes queried in relation to the MAGs generated. Individual assembly statistics are available in Table 1.

Genes associated with denitrification, specifically those associated with nitrate (*nap* and *nar*) and nitrite (*nrf* and *nir*) reduction were considered in MAG analysis as the reduction of oxidized nitrogen species can be utilized in sulfur oxidation reactions. Genes identified as *nap* were more abundant than those putatively identified with homology to *nar* (Figure 4). No MAGs had both genes, and all *nap* genes were found in higher quality MAGs (1, 2, 7, and 9), with the two instances of *nar* genes identified in lower quality MAGs (21 and 29). Nitrite reductase genes were more abundant across MAGs (*nrf* - 1, 2, 7, 12, 21, and 22; *nir* - 1, 3, 5, 8, 20, 24, 25, and 28), however, were not especially abundant. Only two MAGs appear to have the potential to completely reduce nitrate (NO_3_^-^) to ammonium (NH_4_^+^; MAG 1 - *nap, nrf*, and *nir*; MAG 7 - *nap* and *nrf*).

**Figure 4.**
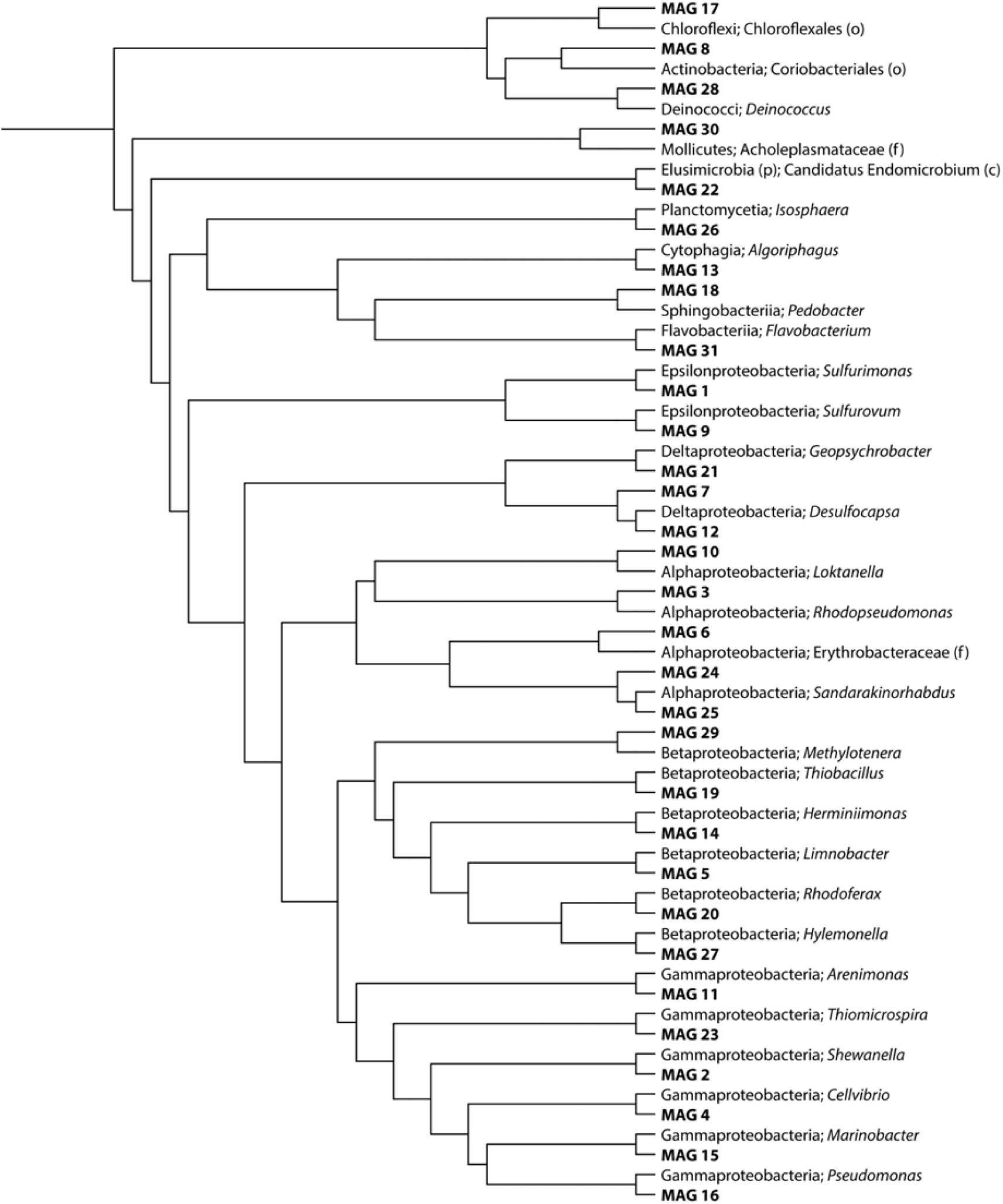
Cladogram of MAGs. Cladogram of metagenome assembled genomes (MAGs) and nearest neighbors to confirm putative taxonomy. Created in the interactive tree of life (iTOL) using the output tree from pparser after initial identification in CheckM.

Cytochrome *c* oxidase (*cbb3*-type) was searched as a marker for aerobic respiration (Pitcher and Watmough, 2004), and hydrogenases *hydAB* (Takai et al., 2005) and *hoxU* (Roalkvam et al., 2015) genes were also investigated due to their importance within *Epsilonproteobacterial* metabolisms. Cytochrome *c* oxidase *cbb3* was found in the majority (18 of 31) MAGs. Only one instance of *hydA* was found in a contig associated with MAG 8. *HoxU* gene abundance was found in 9 out of 31 MAGs (Figure 3b). The gene *hpnp*, a hopanoid biosynthesis gene that codes for an enzyme that methylates hopanoids at the C2 position, is often used as a diagnostic for biomarker potential (Garby et al., 2013). From an astrobiological perspective identifying biomarkers such as hopanoids in extreme environments can be useful in conjunction with potential metabolisms. No *hpnp* (hopanoid synthesis) genes were identified within the BFP MAGs.

### Sulfur Oxidation is Abundant Across BFG Metagenome Assembled Genomes

Of greatest interest for this study were sulfur cycling associated genes (including both the oxidation and reduction of sulfur species). MAGs were searched for well-known genes involved in sulfide oxidation (e.g., *fcc, sqr*), sulfite oxidation (*sorAB*), and thiosulfate oxidation (*sox*). A gene critical to sulfide oxidation, *fcc*, was found in MAGs 1, 5, 9, 19, 20, and 27; two of these (1 and 9) correspond to nearest neighbors in the *Epilonproteobacteria* class (*Sulfurimonas* and *Sulfurovum*). The other five *fcc*-containing MAGs correspond to organisms within the *Betaproteobacteria* (Figure 4). Sulfide-quinone reductase (*sqr*) was only observed in four MAGs (3, 5, 6, and 25), three of which (3, 6, and 25) are *Alphaproteobacteria* and the other most closely related to the genus *Limnobacter* (also containing the aforementioned *fcc* gene). No MAGs showed the presence of genes necessary for sulfite dehydrogenase (*sorAB*).

The Sox system is a widely-studied pathway for the complete oxidation of thiosulfate (S_2_ O_3_^2-^) to sulfate (SO_4_^2-^). This metabolism is typically broken up into four different multi-gene proteins, which make up the larger Sox enzyme complex (*soxAX, soxYZ, soxB*, and *soxCD*). Sox genes were identified in MAGs 1 (*soxXA, soxYZ*, and *soxB*) and 9 (*soxXA* and *soxYZ*), which were identified as most closely related to the genera *Sulfurimonas* and *Sulfurovum* within the *Epsilonproteobacteria*, respectively. Sox genes were also identified in MAGs (5, 10, 14, 15, 19, 20, 22, 23, 27, and 29), with complete operons observed in MAGs 5, 10, 15, 20, and 23 (Figure 3b). All but one of these MAGs are associated with *Proteobacterial* classes, including *Betaproteobacteria* (5, 14, 19, 20, 27, and 29), *Alphaproteobacteria* (10), and *Gammaproteobacteria* (15 and 23), with MAG 22 being nearest neighbors to the candidate class *Endomicrobium* (Phylum *Elusimicrobia*). Not all MAGs had full sox gene complexes (Figure 3b) but all had more than one copy of Sox-associated genes (e.g. multiple copies of *soxA;* Figure S3). Five MAGs (5, 10, 15, 20, and 23) have genes present for all four of the sox-related proteins (Figure 3b).

Three other sulfur-oxidizing genes that co-associate are *apr, sat*, and *qmo*, which carry out the oxidation of sulfite to sulfate (Frigaard and Dahl, 2008). In the green sulfur bacteria these genes are seen as part of the *sat*-*aprBA*-*qmoABC* gene operon; however, a similar *aprAB-qmoABC* operon is also present in sulfate-reducing bacteria (Dahl and Friedrich, 2008). The *sat*-*aprBA*-*qmoABC* operon was identified in MAG 19, (genus *Thiobacillus*). MAG 7, putatively identified as *Desulfocapsa*, contains both *apr* and *qmo* genes; however, this is not the case in MAG 12, also identified as *Desulfocapsa*, where *apr* is absent, but *sat* is present alongside *qmo*. Overall, only three MAGs had *qmo* genes present (MAGs 7, 12, and 19) out of the 31 queried. In only one instance in the BFP MAGs were *apr* and *sat* found together (MAG 5), which is most closely related to the *Betaproteobacterial* genus *Limnobacter*. Otherwise, *apr* (found in four MAGs) was detected less often than *sat* (present in 15 MAGs), which can be seen in Figure 3b. However, gene count totals were comparable with *apr* having 16 and *sat* having 26 instances, respectively (Figure 5), most of which can be attributed to MAG 19, where 9 instances of *apr* were present. The 3’-phosphoadenosine 5’-phosphosulfate (PAPS) gene, often found as an intermediate in dissimilatory sulfide oxidation (Han and Perner, 2015), was also identified in the majority of MAGs (18 of 31).

**Figure 5.**
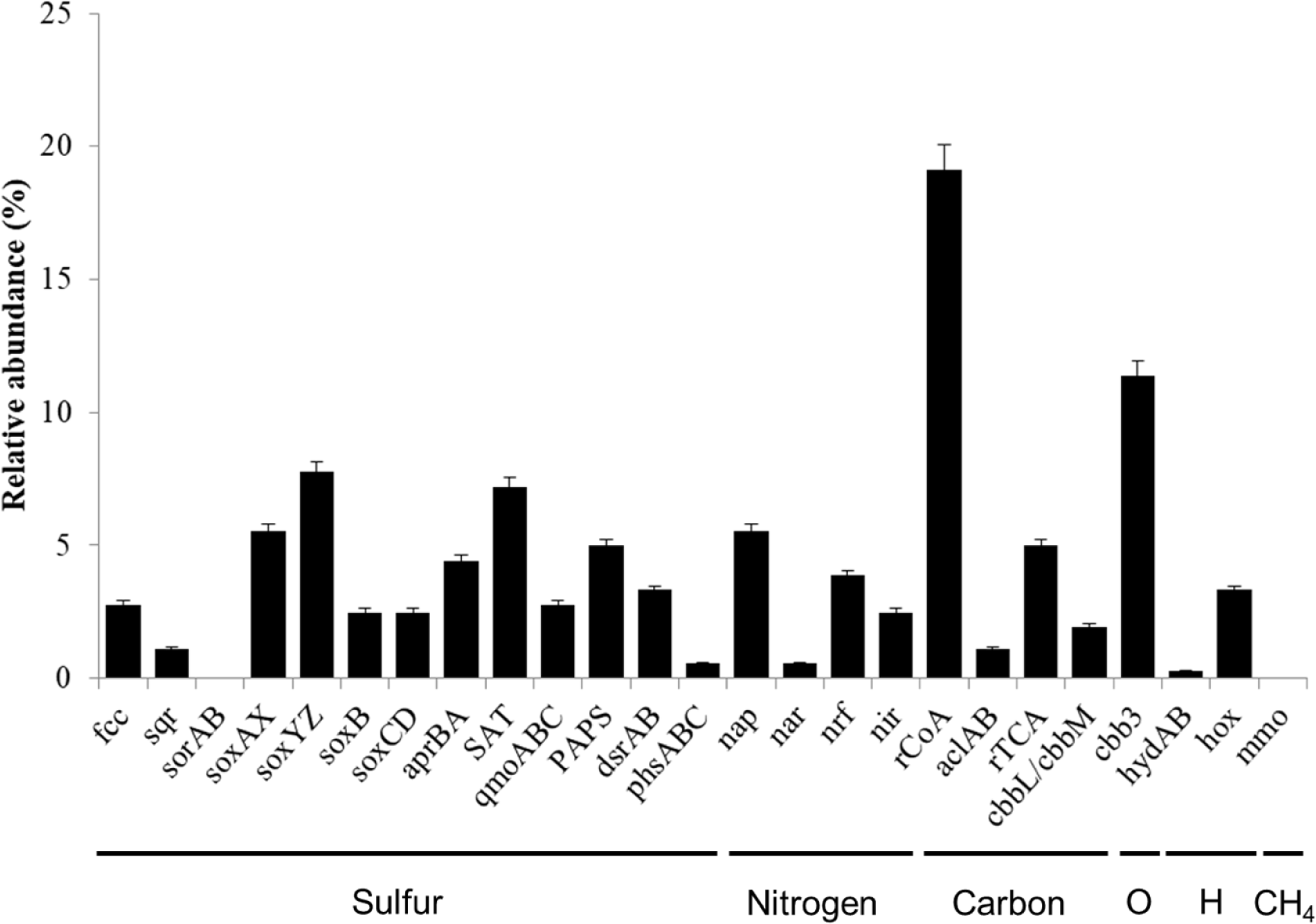
Observed gene sequences within queried MAG contigs. Total number of select metabolism-associated gene sequences found when queried against all 31 reportable MAGs. Gene numbers are normalized to the total number of genes identified within the co-assembly MAGs. *fcc*, sulfide dehydrogenase; *sqr*, sulfide-quinone reductase; *sorAB*, sulfite dehydrogenase; soxXA, soxYZ, soxB, soxCD, proteins making up the Sox enzyme complex; *aprBA*, SAT, *qmoABC*, gene operon; PAPS, 3’-phosphoadenosine 5’-phosphosulfate; *dsrAB*, dissimilatory sulfite reductase; *phsABC*, thiosulfate reductase; *nap*, periplasmic nitrate reductase; *nar*, nitrate reductase; *nrf*, nitrite reductase; *nir*, nitrite reductase; rCoA, reductive acetyl-coenzyme A (Wood-Ljungdahl); *aclAB*, ATP citrate lyase; rTCA, phosphoenolpyruvate carboxylase; *cbbL/cbbM*, ribulose 1,5-bisphosphate carboxylase/oxygenase; *cbb3*, cytochrome *c* oxidase; *hydAB*, hydrogenase I; *hox*, NiFe hydrogenase; *mmo*, methane monooxygenase.

Dissimilatory sulfite-reductase (*dsrAB*) was also queried to identify the respiratory sulfate reduction potential of the system. *dsrAB* was present in three MAGs (7, 12, and 19) two of which correspond to the *Deltaproteobacterial* genus *Desulfocapsa*, while the other appears to be nearest to the *Betaproteobacterial* genus *Thiobacillus*. All three MAGs also have the DsrMKJOP transmembrane gene complex present which often interacts with *dsrAB*, as well as the dissimilatory sulfite-reductase gamma subunit and some form of the assimilatory gene pathway. Two copies of *phs* (thiosulfate reductase), a gene associated with sulfur disproportionation, were identified in MAG 2 (Figures 3 & 5).

## 3 Discussion

Arctic and Antarctic freshwater ecosystems provide insight into low-temperature geochemical sulfur cycling and microbial community dynamics. One such low-temperature environment in the Arctic is Borup Fiord Pass (BFP), a glacial ecosystem dominated by sulfur-rich ice and mineral precipitates. The bioenergetic capability of microbes living in a low temperature Arctic sediment mound at BFP was previously identified through metagenomic sequencing (Wright et al., 2013); however, the work presented increases our understanding of low-temperature microbial metabolisms by expanding the number of sites and sample types beyond surface precipitates by including aufeis samples and fluid from an active spring in 2016. Through the use of metagenomic sequencing we have identified microbial community members from a variety of sample types at BFP that have an abundance of genes related to the oxidation of reduced sulfur species via the *sox* operon including taxa known to perform this process (e.g. *Epsilonproteobacteria – Sulfurimonas* and *Sulfurovum*), as well as others not previously known to carry out this function in these environments (e.g. *Limnobacter* sp.). We conclude that sulfur oxidation may be functionally redundant and more widespread phylogenetically in sulfur-rich polar environments.

In addition to the cycling of sulfur identified at BFP we also searched for core metabolic processes including carbon and nitrogen metabolism. Low levels of nitrate were previously reported at BFP (0.0030 mM in 2009; Wright et al., 2013, and 0.0908 mM in 2014; Lau et al., 2017) within both spring and melt pool fluids, respectively, possibly due to microbial nitrate reduction. Based on gene presence/absence we infer that some MAGs from BFP may be capable of nitrate/nitrite reduction. The *nap* gene, or periplasmic nitrate reductase, is the most abundant identified nitrate reductase (Figure 5), with multiple copies identified in MAGs 1, 2, and 9 (*Sulfurimonas, Shewanella*, and *Sulfurovum*, respectively). This suggests that sulfur oxidizing microorganisms (SOMs) such as *Sulfurimonas* and *Sulfurovum* found in abundance at some sample sites (e.g., M2 and M4b, respectively; Figure 3a) are capable of oxidizing reduced sulfur compounds and utilizing nitrate as an electron acceptor. The *nir* gene, a nitrite reductase, which reduces nitrite to nitric oxide (NO) is slightly less abundant than *nrf*, which reduces nitrite to ammonium (NH_4_^+^). It should be noted that only MAGs which show the presence of *nap* also have co-occurrence of the *nir* gene, which suggests that denitrification in the system may occur via nitric oxide and/or nitrite. While the majority organisms at BFP are most likely using oxygen as an electron acceptor, metagenomic evidence supports the idea that anaerobic denitrification is a viable mechanism of low-temperature respiration.

The utilization of carbon and how it plays a part in microbial growth dynamics at BFP is important to determine, particularly how organic (or fixed) carbon might enter the system. Organic carbon is available within the BFP system (Lau et al., 2017) and has a potential source from nearby shale units (Grasby et al., 2003), however, previous research by Wright, et al. (2013) indicated that the most likely form of energy production in the system is via aerobic oxidation of S_0_ rather than that of organic carbon. Because of this, sulfide oxidation coupled with CO_2_ fixation is potentially one of the most important metabolisms present at BFP. One such phylum containing organisms capable of autotrophy and sulfur oxidation are the *Epsilonproteobacteria* (Campbell et al., 2006). Genes related to one such carbon fixation pathway, the Wood-Ljungdahl pathway (reductive acetyl-coenzyme A pathway, rCoA), appear across all but three of the MAGs (14, 22, and 31). This is in contrast to genes related to the reductive TCA (rTCA) cycle that were present in 17 out of 31 MAGs (including *acl* which was found in two MAGs). The rTCA cycle has previously been implicated as the primary means of converting inorganic carbon into biomass for organisms within the *Epsilonproteobacteria* (Dahl and Friedrich, 2008; Hügler et al., 2010). However, the reductive acetyl-CoA cycle is the predominant carbon fixation pathway found throughout all of our MAG data. Similarly, Momper et al. (2017) found the reductive acetyl-CoA pathway to be the preferred pathway most likely linked to energy limitations within deep subsurface sediments. However, while Borup Fiord Pass (BFP) is an extreme surface environment, organic carbon data and bioenergetics calculations suggest that BFP microbes are not at the lower thermodynamic limit of life (Wright et al., 2013; Lau et al., 2017). Additionally, the reductive acetyl-CoA pathway requires anoxic conditions, which is contrary to the oxygen-rich surface sampling conditions at BFP. A possible explanation for reductive acetyl-CoA carbon fixation at BFP might be the establishment of anoxic zones just beneath the surface of any mineral precipitate or melt pool sediment materials. Within glacial cryoconite sediments similar to those sampled, an anoxic zone can re-establish at 2 to 4 mm from the surface of the sediment within an hour after disturbance (Poniecka et al. 2017).

It should be noted that aerobic metabolism genes are also present in abundance across sample types at BFP as evidenced by the *cbb3* cytochrome *c* oxidase gene. Cytochrome *c* oxidases have been shown to operate under low oxygen tension (Koh et al., 2017) and could still be useful in hypoxic zones near surface sediments; however, this has not yet been established for the *cbb3* cytochrome oxidase like those identified here. Sulfur oxidation and carbon fixation may occur under micro-oxic conditions where the reductive acetyl-CoA pathway is still effective as the dominant carbon fixation process.

The research presented here identified sulfur-oxidizing genes in MAGs from different sites and site types at BFP, and across multiple different taxonomic lineages. Previous research at BFP identified (from both spring fluid and mineral deposits) a number of the same organisms found within our identified MAGs, including: *Marinobacter, Loktanella* (Grasby et al., 2003; Gleeson et al., 2011), and the sulfur oxidizers *Sulfurimonas, Sulfurovum*, and *Sulfuricurvum* (Gleeson et al., 2011, 2012; Wright et al., 2013; Trivedi et al., 2018). Our work has revealed that MAGs belonging to previously identified genera likely oxidize reduced sulfur compounds in addition to organisms that were not previously predicted to be taking part in such metabolic processes. One MAG (MAG 5), putatively identified within the genus *Limnobacter*, contains genes associated with the oxidation of hydrogen sulfide (*fcc, sqr*), as well as a complete pathway (sox) needed to oxidize hydrogen sulfide to sulfate. Chen et al. (2016) reported the full sox pathway within a *Limnobacter* sp. in an anaerobic methane oxidizing microbial community, while other studies have reported its ability to oxidize thiosulfate to sulfate (Kämpfer et al., 2001; Lu et al., 2011). However, we are unaware of any previous research linking *Limnobacter* sp. to environmental sulfide oxidation.

*Loktanella* sp. were also previously identified at BFP in 16S rRNA gene clone libraries (Gleeson et al., 2011) as well as in other low-temperature environments throughout the Canadian Arctic (Perreault et al., 2007; Niederberger et al., 2009; Battler et al., 2013). *Loktanella* was previously reported as containing *soxB* from samples obtained at Gypsum Hill, a perennial Arctic spring on Axel Heiberg Island (Perreault et al., 2008). The Perreault, et al. study also found that the *Loktanella* sp. was part of a consortium along with a *Marinobacter* sp. similar to what was found in previous samples at BFP (Gleeson et al., 2011; Trivedi et al., 2018).

In contrast to previous 16S rRNA gene sequencing analyses, we putatively identified the genera *Herminiimonas* and *Rhodoferax* within the class *Betaproteobacteria* through whole genome sequencing and binning. Despite its identification within a 120,000-year-old Greenland ice core (Loveland-Curtze et al., 2009), *Herminiimonas* is rarely identified within polar environments. Moreover, the data on the capability of *Herminiimonas* to oxidize sulfur is equally sparse. In a recent study, Koh et al. (2017) found that *Herminiimonas arsenitoxidans*, which was found to oxidize arsenite, was able to oxidize sulfur when grown on trypticase soy agar. Conversely, *Rhodoferax* (also known as purple non-sulfur bacteria), another *Betaproteobacterium* that is commonly found in polar environments (Madigan et al., 2000; Lutz et al., 2015), are facultatively photoheterotrophic; however, some are capable of photoautotrophy when sulfide is used as an electron donor (Ghosh and Dam, 2009). A BFP MAG putatively identified as *Rhodoferax* (MAG 20) also shows the presence of genes related to the reductive acetyl-CoA pathway, which could confirm that this particular organism is capable of autotrophy and may serve as a primary producer in this sulfur-dominated polar ecosystem.

The soxCD complex has been shown to be responsible for the oxidation of sulfur to thiosulfate, and in organisms that lack this complex the sulfur is either stored inside the cell or excreted (Hensen et al., 2006). The lack of soxCD in BFP MAGs (e.g. MAGs 1, 9, and 19) from known sulfur-oxidizing microorganisms (*Sulfurimonas, Sulfurovum*, and *Thiobacillus*) is an interesting finding. In fact, this may be one possible biological explanation for the abundance of elemental sulfur precipitated across the surface of BFP. Some organisms may employ other pathways like reverse dissimilatory sulfite reduction (rDsr) to oxidize accumulations of S^0^ (Hamilton et al., 2015); however, the low prevalence of any sulfite reductase genes in the BFP system could point to an accumulation of elemental sulfur rather than additional biogenic oxidation. Thermodynamically sulfur oxidation should occur abiotically, however, at such low temperatures, this abiotic process may be kinetically slow without biological catalysis.

In addition to sulfur oxidation, dissimilatory sulfate reduction is also a key component in the sulfur cycle and potentially in energy generation at BFP. Out of the 31 identified MAGs, three (MAG 7, 12, and 19) contain dissimilatory sulfite reductase (*dsrAB*) genes, a key step in the complete reduction of sulfate to sulfide. Two MAGs, MAG 7 and 12, are most closely related to the genus *Desulfocapsa*, a known sulfur disproportionator within the *Deltaproteobacteria* (Finster et al., 1998, 2013). This genus was previously reported at BFP via 16S rRNA gene sequencing (Trivedi et al., 2018). It should also be noted that while genes specific to disproportionation (*phs*) were searched for, only one instance was found, within MAG 2, which is most closely related to the genus *Shewanella*. Another MAG most closely related to the *Betaproteobacterial* genus *Thiobacillus* (MAG 19) also contains a complete *dsr* operon. Interestingly, MAG 19 is the only MAG that appears capable of both sulfur-oxidation via the sox pathway and sulfate-reduction via dissimilatory sulfite reductase. *Thiobacillus* sp. are traditionally known as sulfur oxidizing microorganisms (SOMs) and have been found in polar environments previously (Boyd et al., 2014; Purcell et al., 2014; Harrold et al., 2016); however, it has also been suggested that certain *Thiobacillus* species carry out reverse dissimilatory sulfite reduction (rDsr, or reverse dsr) as an alternative means to oxidize reduced sulfur compounds (Loy et al., 2009; Purcell et al., 2014). This could well be an alternative pathway that is present in the BFP system; however, it would require transcriptomic evidence to confirm its activity. It should be noted that MAG 19 appears to lack soxCD; however, the MAG is only ∼73% complete, which may indicate that its full metabolic potential is not represented.

The abundance of sox genes present in BFP across sample types and multiple bacterial lineages suggests that sulfur cycling is functionally redundant in the BFP ecosystem. Microorganisms in cold, deep subsea floor aquifers have also been identified with high degrees of functional redundancy (Tully et al., 2018). Functional redundancy was recently identified as a commonplace process in environmental systems by Louca et al. (2018). The authors speculated that the degree of functional redundancy in an ecosystem is largely determined by the type of environment, in contrast to the belief that species should inhabit distinct microbial niches independent of environment. Moreover, Louca et al. call into question a previous idea that functional redundancy in an ecosystem might infer a form of ‘neutral co-existence’ between competing microorganisms (Loreau, 2004). However, their findings challenge this, as coexisting microorganisms will often differ in ways that will affect how they grow under environmental conditions. Based on these ideas it is highly probable that functional redundancy may be an important survival mechanism in use by microorganisms at BFP.

At BFP a core microbial community not predicted to participate in sulfur cycling exists as identified by a 16S rRNA gene sequencing (Trivedi et al., 2018). Instead our metagenomic data indicate that multiple phylogenetically disparate core communities may have the potential to utilize sulfur compounds for growth. As suggested by Louca et. al (2018), coexisting organisms may not be in equal abundance (such as those identified within the core community) as there will always be differences in functional ability based on enzyme efficiencies and growth kinetics. This also supports previous research of the BFP system, where “blooms” of SOMs over short time periods occurred, followed by a return to a basal microbial community (Trivedi et al., 2018). Such blooms of SOMs could be facilitated from above by the input of new microbiota via aeolian transport in meteorologic storm events whereby snow storms contain significant amounts of microbiota and genetic potential (Honeyman et al., 2018). If there is a large influx of reduced sulfur compounds from beneath the ice, some phyla, like the occasionally (e.g. conditionally) rare *Epsilonproteobacteria* (Trivedi et al., 2018), may be preferentially able to use these compounds, much like in deep sea vent fluids (Akerman et al., 2013) leading to an increase in their relative abundance until settling to previously identified levels. Undoubtedly, sulfur oxidation is the predominant and preferred metabolism in these sulfur-rich surface sites at BFP.

The response to ecological change by conditionally rare taxa (i.e., the low representation of microbiota in any environment that can become dominant upon environmental condition change) thus reflects a biological manifestation of functional redundancy. The presence of functional redundancy found at BFP is due to the multiple environmental pressures at the site such as extreme cold, and high concentrations of multiple sulfur species (e.g., H_2_S, HS^-^, S^0^, S_2_ O_3_^2-^, and SO_4_^2-^) geochemically produced in the subsurface (Wright et al., 2013). These compounds then emanate to the surface where microorganisms then take advantage of the abundance of reduced sulfur for growth. This may be driven by the unique convergence of lower levels of organic carbon, low temperatures, and an abundance of reduced sulfur compounds at BFP, which may not necessarily be the case in all highly sulfidic systems.

These results show that in low-temperature environments, functional redundancy of key metabolisms is an active strategy for multiple bacterial lineages to coexist. The results presented here reflect the potential metabolisms present at BFP generated from whole genome sequencing and the production of MAGs. The next step in truly defining how microorganisms are able to survive and thrive in the BFP system will be to undertake metatranscriptomic sequencing. This next step in the genomic characterization of this ecosystem will allow us to differentiate between the metabolic potential and active processes within the system. Our data show that sulfur oxidation is functionally redundant at BFP and may be utilized by more microorganisms across the bacterial domain than would be expected.

## 4 Materials and Methods

### Sample Collection

BFP sample material was collected during multiple days between 21 June and 2 July 2014, 4 July 2016, and 7 July 2017. Samples included: 1) sedimented sulfur-like cryoconite material and fluid found in glacial surface melt pools (labelled M2 & M4b); 2) filtered fluid from thawed aufeis (labelled A4b & A6); 3) aufeis surface precipitate samples (material scraped from on top of the aufeis) from 2017 (labelled 14C, 3B, 4E, & 6B); and 4) spring fluid from 2016 (labelled BFP16 Spring); see Table S1 and Figures 3a & S2). Cryoconite sediment, aufeis, and spring fluid samples for DNA extraction were filtered through 0.22 µm Luer-lok Sterivex™ filters (EMD Millipore; Darmstadt, Germany), which were then capped and kept on ice in the field until returning to the laboratory where they were stored at −20 °C until extraction. Aufeis surface scrapings (mineral precipitate samples) from the 2017 field campaign were collected using sterile and field-washed transfer pipettes and up to 0.5 g of material were mixed in ZR BashingBead™ lysis tubes containing 750 ml of DNA/RNA Shield™ (Zymo Research Corp.). Samples were shaken and kept at 4 °C until returned to the laboratory, where they were then stored at −80 °C until DNA/RNA extraction could be performed.

### DNA Extraction and Metagenomic Library Preparation

DNA extraction of 2014 melt pool, aufeis, and 2016 spring fluid samples was performed using the PowerWater® Sterivex™ DNA Isolation Kit (MO BIO Laboratories, USA), and DNA extraction of 2017 surface precipitate samples was extracted using the ZymoBIOMICS™ DNA/RNA Mini Kit (Zymo Research Corp.). All extractions were performed according to the manufacturer’s instructions. Extracted DNA concentrations were determined using a QuBit 2.0 fluorometer (Thermo Fisher Scientific, Chino, CA, USA) and the QuBit dsDNA HS assay (Life Technologies, Carlsbad, CA, USA).

Genomic DNA was first cleaned using KAPA Pure Beads (KAPA Biosystems Inc., Wilmington, MA, USA) at a final concentration of 0.8X v/v prior to library preparation. Two metagenomic sequencing runs were completed for the included sample set. The first (including samples BFP14 A4b, A6, and M4b) were normalized to 1 ng of DNA in 35 µl of molecular biology grade water, and prepared using the KAPA HyperPlus Kit (KAPA Biosystems Inc., Wilmington, MA, USA) and KAPA WGS adapters. Fragmentation and adapter ligation was performed according to the manufacturer’s instructions. Prepped samples were quality checked on an Agilent 2100 Bioanalyzer (Agilent Genomics, Santa Clara, CA, USA) and submitted to the Duke Center for Genomic and Computational Biology (Duke University, Durham, NC, USA). Final pooled libraries were run on an Illumina HiSeq 4000 (Illumina, San Diego, CA, USA) using PE150 chemistry. The second sequencing run (which included samples BFP14 M2, BFP16 Spring, and BFP17 3B, 4E, 6B, and 14C) was prepped using the Nextera XT DNA Library Preparation Kit (Illumina, San Diego, CA, USA) in conjunction with Nextera XT V2 Indexes. Libraries were hand normalized and quality checked on an Agilent 2100 Bioanalyzer (Agilent Genomics, Santa Clara, CA, USA). Samples were submitted to the Duke Center for Genomic and Computational Biology (Duke University, Durham, NC, USA). These libraries were sequenced on an Illumina HiSeq 2500 Rapid Run using PE250 chemistry.

### Metagenomic Sequencing Assembly and Analysis

Processing of metagenomic sequencing was performed on the Summit High Performance Cluster (HPC) located at the University of Colorado, Boulder (Boulder, CO, USA). Assembly of whole-genome sequencing reads used a modified Joint Genome Institute (JGI; Walnut Creek, CA, USA) metagenomics workflow. We began by using BBDuk of the BBMap/BBTools (v37.56; Bushnell, 2014) suite to trim out Illumina adapters using the built-in BBMap adapters database. Trimmed sequences were checked for quality using FastQC v11.5 (Andrews, 2014) to ensure that adapters were removed successfully. Trimmed paired-end sequences were then joined using the PEAR package (Zhang et al., 2014) v0.9.10 using a p-value of 0.001. Adapter trimmed files, both forward (R1) and reverse (R2) were then processed using BBTools ‘repair.sh’ to ensure read pairs were in the correct order. This script is designed to reorder paired reads that have become disorganized, which can often occur during trimming. The BFC (Bloom Filter Correction) software tool was then used to error correct paired sequence reads (Li, 2015). Reads were again processed with BBTools ‘repair.sh’ to ensure proper read pairing and order. Next, Megahit v1.1.2 (Li et al., 2015) was used to generate both a co-assembly, as well as individual sample assemblies from the quality controlled reads. Post assembly, the co-assembly was processed with ‘anvi-script-reformat-fasta’ for input into Anvi’o, and contigs below 2500 bp were removed. Bowtie2 v2.3.0 (Langmead and Salzberg, 2012) was then used to map individual sample sequence reads onto the co-assembly, needed for the Anvi’o pipeline. Anvi’o v4 (Eren et al., 2015) was then used to characterize, and then bin co-assembled contigs into metagenome assembled genomes (MAGs). The taxonomic classifier Centrifuge v1.0.2-beta (Kim et al., 2016) was used to provide a preliminary taxonomy of the co-assembly contigs, useful for Anvi’o binning. The NCBI Clusters of Orthologous Groups (COGs; (Tatusov et al., 2000; Galperin et al., 2015)) database was used within Anvi’o to predict COGs present within the co-assembly, and CONCOCT v1.0 (Alneberg et al., 2014) was employed to create genome bins using a combination of sequence coverage and composition of our assembled contigs. Anvi’o was then used to visualize the putative bins in conjunction with other metadata to refine and identify MAGs with 50% completeness and 10% redundancy. The scripts used in the processing of these data (up to the point of Megahit-generated contigs) are publicly available at https://doi.org/10.5281/zenodo.1302787.

Anvi’o refined bins were then processed using CheckM v1.0.11 (Parks et al., 2015) where a total of 31 bins were selected as MAGs (Metagenome Assembled Genomes) that followed the > 50% completeness and < 10% redundancy criteria. MAGs that met these criteria were then renamed; from 1-31 in the order of highest completeness to lowest (Table 1). CheckM relies on other software including pplacer v1.1.alpha17 (Matsen et al., 2010) for phylogenetic tree placement, prodigal v2.6.3 (Hyatt et al., 2012) for gene translation and initiation site prediction, and HMMER v3.1b2 (Eddy, 2011) which analyzes sequence data using hidden Markov models. The pplacer generated tree was then visualized using the interactive Tree of Life (iTOL; (Letunic and Bork, 2016)) to search for nearest neighbors of identified MAGs (see Figure 4). Finally, to confirm general taxonomic classification and annotate genes within our MAGs, samples were processed using the Rapid Annotations using Subsystem Technologies (RAST) server (Aziz et al., 2008). Genes of interest to this study were identified within the RAST web interface. Further taxonomic classification was done for MAGs 7 and 12 using the online tool JSpeciesWS (Richter et al., 2016), which uses pairwise genome comparisons to measure the probability of two or more genomes. Tests within JSpeciesWS include average nucleotide identity (ANI) via BLAST and MUMmer and correlation between Tetra-nucleotide signatures. Average nucleotide identity (ANI) was used to determine if MAGs 7 and 12 (putatively classified as *Desulfotalea*) were more closely related to the genus *Desulfocapsa* as was previously identified within BFP samples (Trivedi et al., 2018).

### Data Accessibility

Raw metagenomic sequencing data are available at the NCBI sequencing read archive (SRA) under the accession numbers SRR10066347-SRR10066355 (https://www.ncbi.nlm.nih.gov/sra). Assembled MAGs are provided via FigShare under the following DOI: 10.6084/m9.figshare.9767564. Additionally, the bioinformatics workflow up to the point of assembled contigs can be found under DOI: https://doi.org/10.5281/zenodo.1302787.

## 6 Author Contributions

AST, JRS, and SEG led the design of the study. Fieldwork was conducted by CBT, GEL, SEG, AST, and JRS. Laboratory work was performed by CBT and GEL, and bioinformatic analysis was done by CBT and BWS. All authors interpreted results. CBT is the primary author of the manuscript with contributions and guidance from all other authors.

## 8 Acknowledgements

Field logistics and travel to/from Borup Fiord Pass were supported through Natural Resources Canada’s (NRCan) Polar Continental Shelf Program (PCSP). JRS and SEG thank Dr. Karsten Piepjohn and the Federal Institute for Geosciences and Natural Resources at BGR, Hanover, Germany for support during the 2017 field campaign of the Circum-Arctic Structural Events-19 (CASE-19) expedition.

## Supplementary Materials

**Table S1.** Co-assembly metadata. These data represent raw values from Megahit prior to mapping through Bowtie2 and final processing via Anvi’o.

| Sample | Type | Total bp | Contigs | Max contig (bp) | Avg (bp) | N50 (bp) |
| --- | --- | --- | --- | --- | --- | --- |
| BFP14_A4b | Aufeis | 85484280 | 30017 | 380944 | 2848 | 3571 |
| BFP14_A6 | Aufeis | 82535183 | 33359 | 128624 | 2474 | 2871 |
| BFP14_M2 | Melt pool | 10144305 | 4841 | 418178 | 2095 | 1977 |
| BFP14_M4b | Melt pool | 1.59E+08 | 71593 | 111481 | 2215 | 2359 |
| BFP16_Spring | Spring fluid | 60894340 | 18996 | 206795 | 3206 | 4748 |
| BFP17_14C | Aufeis surface | 6.38E+08 | 272035 | 297076 | 2347 | 2611 |
| BFP17_AS3b | Aufeis surface | 1.98E+08 | 78690 | 78857 | 2513 | 3028 |
| BFP17_AS4e | Aufeis surface | 89868307 | 27395 | 44699 | 3280 | 4468 |
| BFP17_AS6b | Aufeis surface | 1E+08 | 34932 | 60991 | 2866 | 4186 |
| **Co-Assembly** |  | **1.16E+09** | **477394** | **406785** | **2440** | **2778** |

**Table S2.** JSpeciesWS results. MAGs 7 and 12 were compared to database genomes for *Desulfocapsa sulfexigens* and *Desulfotalea psychrophila* LSv54. Average nucleotide identity was calculated for BLAST and MUMmer as well as Tetra-nucleotide comparison.

| #ANI BLAST and aligned percentage |  |  |  |  |
| --- | --- | --- | --- | --- |
|  | Desulfotalea psychrophila LSv54 | Desulfocapsa sulfexigens DSM 10523 | MAG 12 | MAG 7 |
| Desulfotalea psychrophila LSv54 | * | 66.86 [20.05] | 67.07 [14.44] | 67.13 [14.59] |
| Desulfocapsa sulfexigens DSM 10523 | 66.75 [18.50] | * | 68.41 [25.07] | 68.16 [24.91] |
| MAG 12 | 67.28 [20.09] | 68.74 [35.44] | * | 81.71 [44.41] |
| MAG 7 | 67.17 [20.68] | 68.48 [36.64] | 82.11 [44.96] | * |

| #ANI MUMmer and aligned percentage |  |  |  |  |
| --- | --- | --- | --- | --- |
|  | Desulfotalea psychrophila LSv54 | Desulfocapsa sulfexigens DSM 10523 | MAG 12 | MAG 7 |
| Desulfotalea psychrophila LSv54 | * | 87.49 [1.04] | 87.97 [0.04] | 85.18 [0.14] |
| Desulfocapsa sulfexigens DSM 10523 | 87.74 [0.47] | * | 82.75 [0.55] | 82.87 [0.26] |
| MAG 12 | 87.97 [0.02] | 82.75 [0.75] | * | 85.30 [33.12] |
| MAG 7 | 85.18 [0.17] | 82.84 [0.41] | 85.30 [34.60] | * |

| #Tetra results |  |  |  |  |
| --- | --- | --- | --- | --- |
|  | Desulfotalea psychrophila LSv54 | Desulfocapsa sulfexigens DSM 10523 | MAG 12 | MAG 7 |
| Desulfotalea psychrophila LSv54 | * | 0.90901 | 0.86371 | 0.84073 |
| Desulfocapsa sulfexigens DSM 10523 | 0.90901 | * | 0.92621 | 0.90605 |
| MAG 12 | 0.86371 | 0.92621 | * | 0.98997 |
| MAG 7 | 0.84073 | 0.90605 | 0.98997 | * |

**Figure S1.**
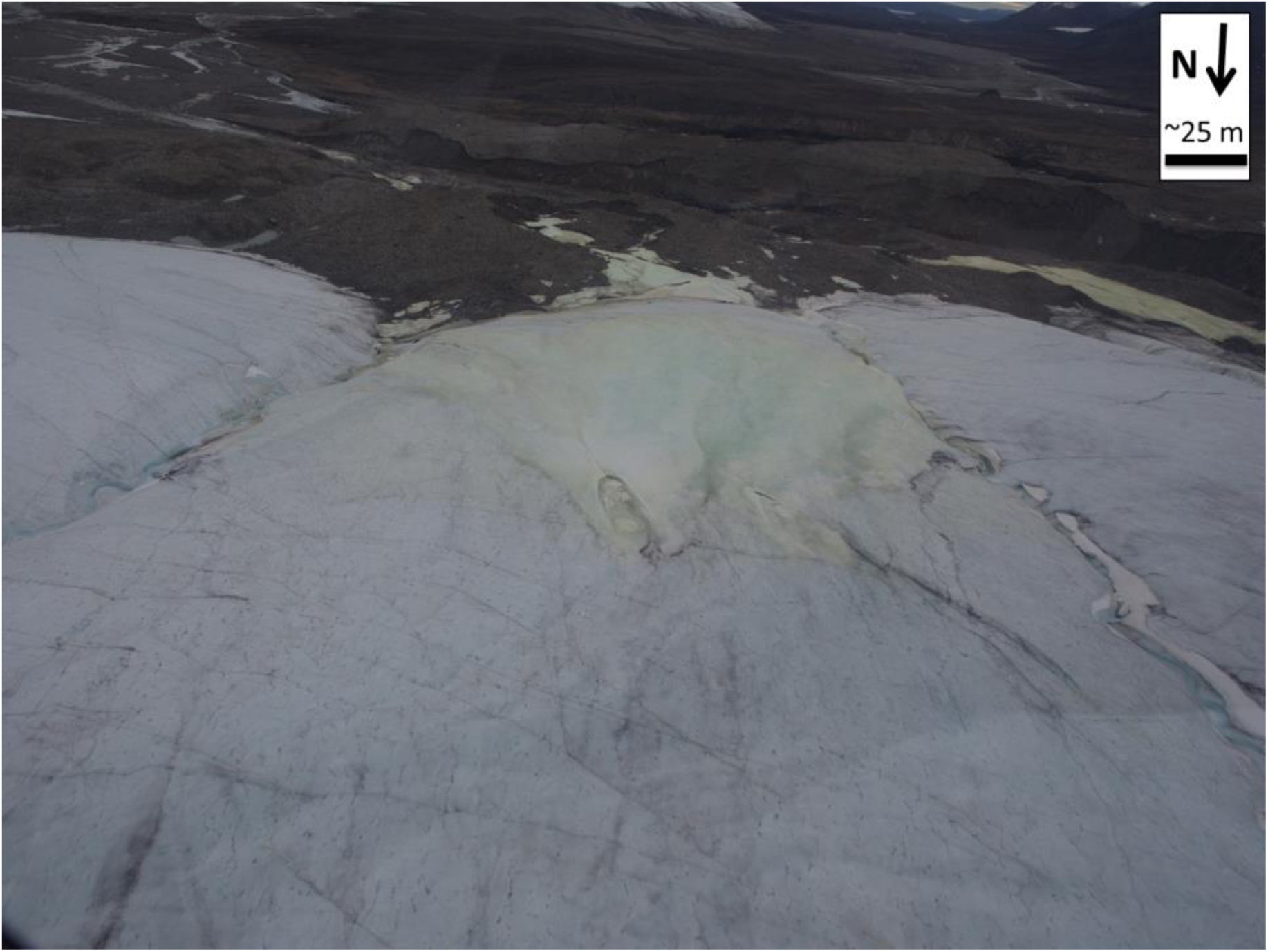
BFP16 Spring location. Shown is the location of the 2016 spring. Its exact location is at the northern-most portion of the yellow-colored area, approximately in the center of the picture. Also shown is the area known as the “Sulfidic Aufeis” from Trivedi, et al. (2018) towards the right edge of the picture. This is the location of sampling from 2014 and 2017 field campaigns.

**Figure S2.**
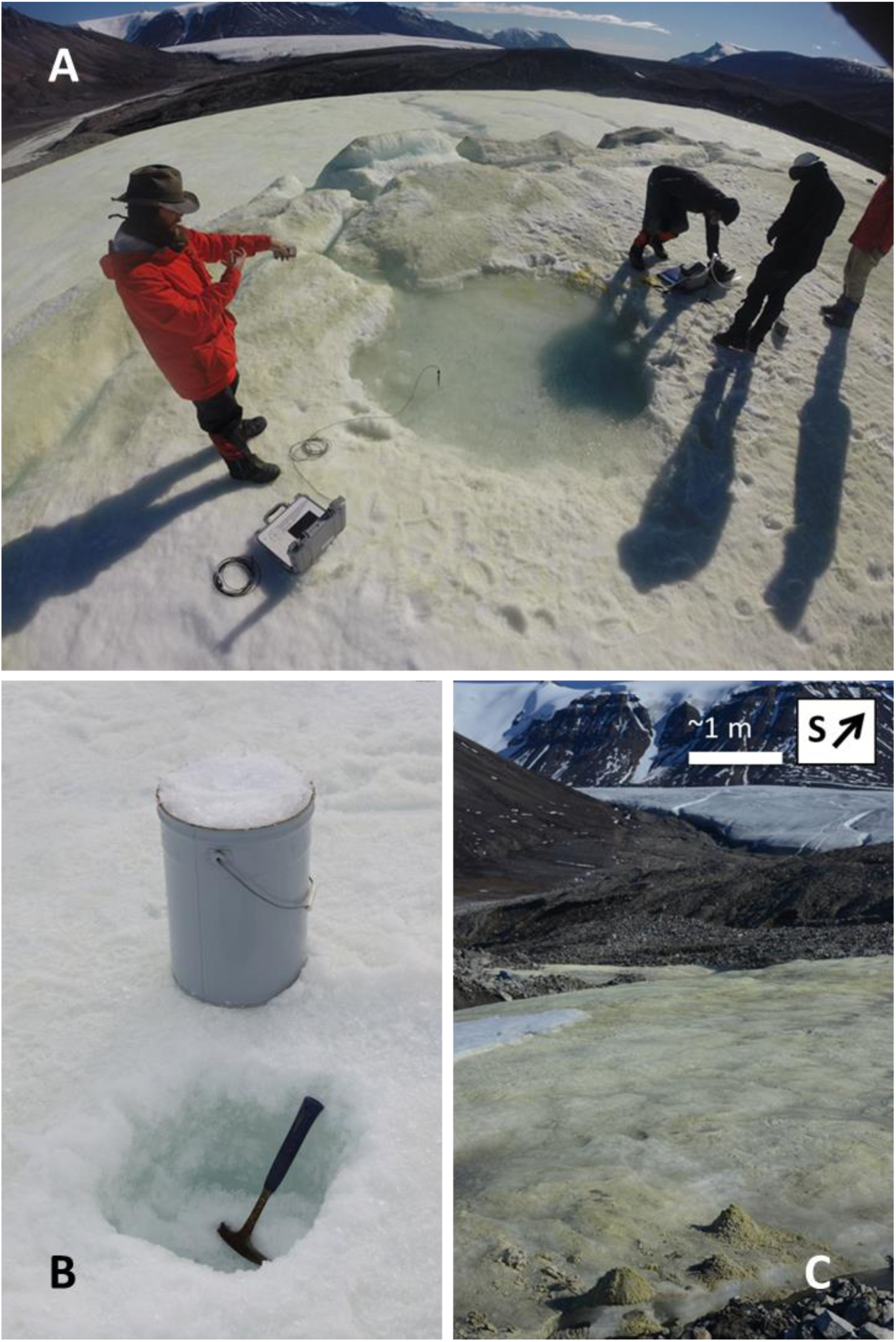
Sample type examples. Examples of sample types used for metagenomes. **A)** Melt pool; **B)** Aufeis sample, where surface ice was removed, a pit dug and ice from this pit was melted and filtered; **C)** Mineral precipitates from the 2017 BFP site.

**Figure S3.**
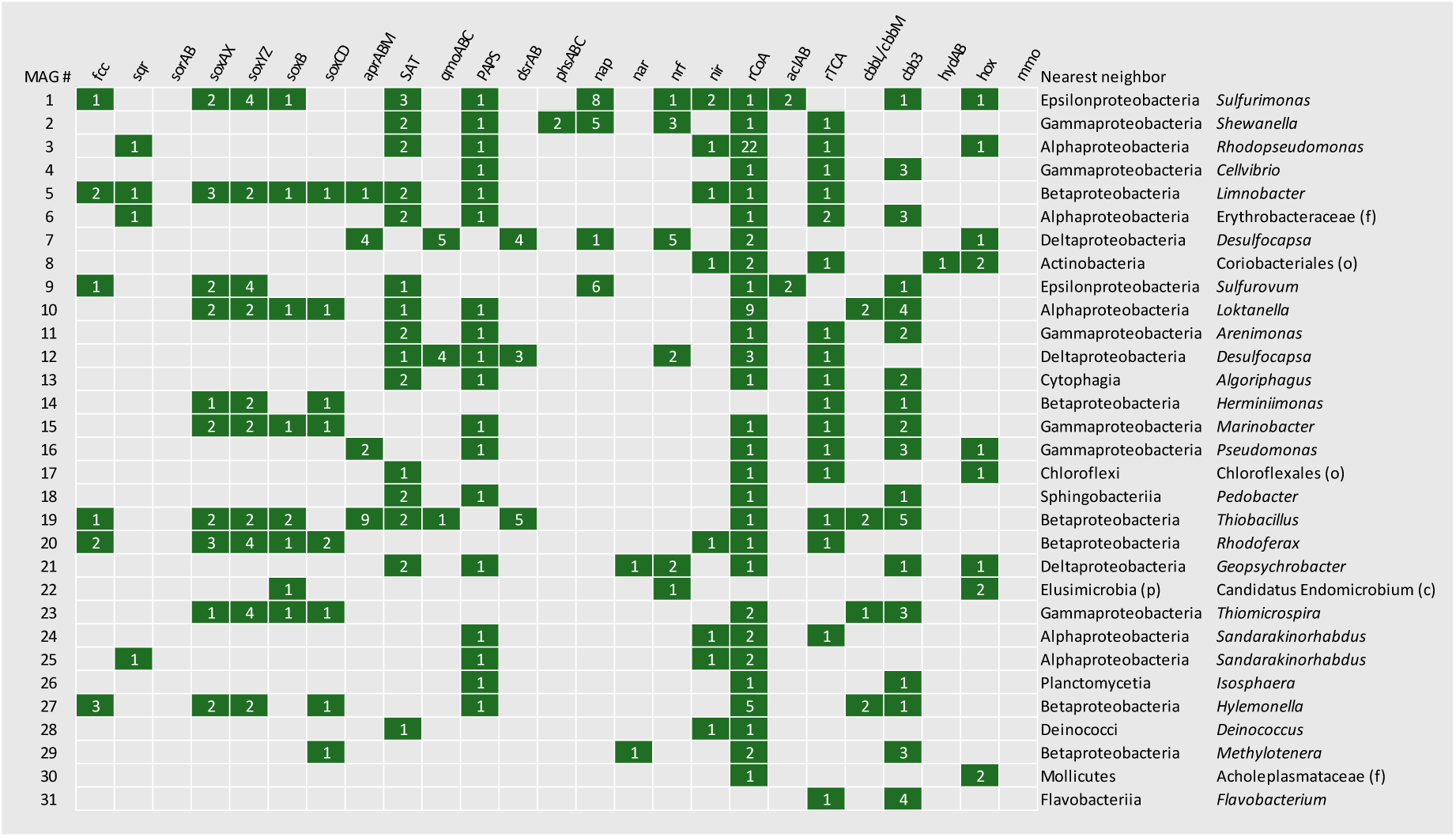
Metabolic gene grid (Figure 3b) with associated counts for each. This grid, presented in the main text as Figure 3b, now with gene counts shown. Each instance of a specific gene found in each MAG was recorded here and then the presence/absence figure was generated. Counts were normalized to generate Figure 5.

